# The *in vitro* and *in vivo* potency of CT-P59 against Delta and its associated variants of SARS-CoV-2

**DOI:** 10.1101/2021.07.23.453472

**Authors:** Dong-Kyun Ryu, Hye-Min Woo, Bobin Kang, Hanmi Noh, Jong-In Kim, Ji-Min Seo, Cheolmin Kim, Minsoo Kim, Jun-Won Kim, Nayoung Kim, Pyeonghwa Jeon, Hansaem Lee, Jeong-Sun Yang, Kyung-Chang Kim, Joo-Yeon Lee, Min-Ho Lee, Sang-Seok Oh, Hyo-Young Chung, Ki-Sung Kwon, Soo-Young Lee

**Author notes:** These authors contributed equally to this work (Dong-Kyun Ryu, Hye-Min Woo). Corresponding authors: Soo-Young Lee, 20, Academy-ro 51 beon-gil, Yeonsu-gu, Incheon, 22014, South Korea.

## Abstract

The Delta variant originally from India is rapidly spreading across the world and causes to resurge infections of SARS-CoV-2. We previously reported that CT-P59 presented its *in vivo* potency against Beta and Gamma variants, despite its reduced activity in cell experiments. Yet, it remains uncertain to exert the antiviral effect of CT-P59 on the Delta and its associated variants (L452R). To tackle this question, we carried out cell tests and animal study. CT-P59 showed reduced antiviral activity but enabled neutralization against Delta, Epsilon, and Kappa variants in cells. In line with *in vitro* results, the mouse challenge experiment with the Delta variant substantiated *in vivo* potency of CT-P59 showing symptom remission and virus abrogation in the respiratory tract. Collectively, cell and animal studies showed that CT-P59 is effective against the Delta variant infection, hinting that CT-P59 has therapeutic potency for patients infected with Delta and its associated variants.

**Highlights:**

- CT-P59 exerts the antiviral effect on authentic Delta, Epsilon and Kappa variants in cell-based experiments.
- CT-P59 showed neutralizing potency against variants including Delta, Epsilon, Kappa, L452R, T478K and P681H pseudovirus variants.
- The administration of clinically relevant dose of CT-P59 showed *in vivo*
- protection against Delta variants in animal challenge experiment.

## 1. Introduction

New lineage B.1.617 of SARS-CoV-2 is considered to cause a steep increase in infection cases and death in India. Concerning transmissibility and pathogenicity, WHO (World Health Organization) designated Kappa (B.1.617.1) as Variants of Interest (VOI) and Delta (B.1.617.2) as Variants of Concern (VOC), respectively. In particular, highly transmissible Delta variants rapidly spread and become dominant in India and other countries [1, 2]. Also, the global dominant Alpha variants were replaced by Delta variants, even in most vaccinated countries such as the United Kingdom and the United States [3, 4].

The B.1.617 variant (L452R/E484Q/Δ681/Δ1072) has three main sub-lineages; B.1.617.1 (E154K/L452R/E484Q/P681R/Q1071H), B.1.617.2 (T19R/G142D/Δ156-157/R158G/L452R/ T478K/P681R/D950N) and B.1.617.3 (T19R/Δ156-157/R158G/L452R/E484K/P681R/ D950N) [5, 6]. The Delta variant has RBD (Receptor Binding Domain) mutations in the spike protein which may contribute to increased infectivity, pathogenicity and immune evasion, resulting in re-infection or resistance to vaccine-elicited antibody and therapeutic antibodies [7-9]. One of Delta-associated variants, B.1.427 (L452R) and B.1.429 (S13I/W152C/L452R) also known as the Epsilon have L452R mutation, causing reduction in antibody neutralization [10-12]. Delta and Kappa variants have a P681R mutation at furin cleavage sites (_681_PRRAR/S_686_). Recent studies showed that P681R enhances spike cleavage and fusion for viral entry, suggesting augmented transmissibility and pathogenicity [13, 14]. In addition, Mexican variant, B.1.1.519 (T478K/P681H/T732A) has common mutation sites with Delta variant at T478 and P681, in common with Alpha variant (Δ69-70/Δ144/N501Y/A570D/ P681H/T716I/S982A/D1118H) [15]. T478K was a previously predominant variant in Mexico and located at the interface of RBD and ACE2 (Angiotensin-Converting Enzyme 2) interaction, thereby potentially impacting viral infection [16]. Thus, it is important to assess the impact of mutations within the spike on COVID-19 therapeutics. The CT-P59 human monoclonal antibody (Regdanvimab) specifically binds to spike of SARS-CoV-2, blocking interaction with ACE2 for viral entry. We have demonstrated that CT-P59 has high potency against the original SARS-CoV-2 in *in vitro* and *in vivo* studies and also showed *in vitro* and *in vivo* potency against B.1.351 and P.1 [17-19]. However, it remains uncertain whether CT-P59 has the therapeutic effect on Delta and its associated variants. To this end, we examined the sensitivity to monoclonal CT-P59 against Delta, Epsilon and Kappa variants by cell assays (plaque reduction neutralization test and pseudovirus assay), and animal test using Delta-infected mice.

## 2. Materials and methods

### 2.1. Viruses

Clinical isolates, Delta (hCoV-19/Korea/KDCA5439/2021), Epsilon (hCoV-19/Korea/KDCA1792/2020, and hCoV-19/Korea/KDCA1793/2020), Kappa (hCoV-19/Korea/KDCA2950/2021) were obtained through National Culture Collection for Pathogens. Pseudoviruses for Delta (L452R/T478K/P681R), Epsilon (S13I/W152C/L452R), Kappa (L452R/E484Q/P681R), L452R, T478K, and P681H were produced as previously described [18].

### 2.2. Biolayer interferometry (BLI)

Binding affinity of CT-P59 to wild type and mutant SARS-CoV-2 RBDs were evaluated by the Octet QK^e^ system (ForteBio). Mutant RBDs (L452R/T478K, and L452R) were purchased from Sino Biological [17, 19].

### 2.3. Plaque Reduction Neutralization Test (PRNT)

PRNT for authentic SARS-CoV-2 variants were carried out as previously described [19]. Briefly, pre-incubated mixture of variant and CT-P59 was inoculated to VeroE6 cells. After incubation, the neutralization of CT-P59 against each variant was determined as the half-maximal inhibitory concentration (IC_50_) using GraphPad Prism6 software.

### 2.4. Pseudovirus assay

Luciferase-based pseudovirus assay was performed as previously described [18]. In brief, Mutant pseudoviruses were generated by PCR-based mutagenesis and produced by transiently transfection. Serially diluted antibodies mixed with pseudoviruses. The mixture inoculated ACE2-HEK293T cells, and incubated for 72 h. Neutralization was assessed by luciferase measurement and IC_50_ calculation

### 2.5. Animal experiments

8-week-old female human ACE2 transgenic mice, tg(K18-ACE2)2Prlmn, were purchased from The Jackson Laboratory. Mice were housed in a certified A/BSL-3 facility at the KDCA. All procedures were approved by the Institutional Animal Care and Use Committee at KDCA (No. KDCA-IACUC-21-026).

### 2.6. Mouse study

For intranasal infection, mice (n=8 for virus only group as a placebo group and n=9/group for each CT-P59 treatment group) were humanely euthanized by intraperitoneal (IP) injection with a mix of 30 mg/kg Zoletil® (Virbac) and 10 mg/kg Rompun® (Bayer). Subsequently, mice were intranasally inoculated with 1×10^4^ PFU of Delta variant in a 30 µL volume. CT-P59 was administered via a single IP injection at dose levels of 5, 20, 40 and 80 mg/kg (equivalent to human doses of 2.5, 10, 20 and 40 mg/kg, based on exposure to mAb from the PK studies) at 8 h post-inoculation. The negative control (n=5) and virus only group mice were administered with formulation buffer via the same route. The body weight and mortality were monitored daily until 6 dpi (days post-inoculation) and lung tissues were collected at scheduled necropsy at 3 dpi (n=3 for the negative control group, n=4 for the virus only group and n=5 for each treatment group) and 6 dpi (n=2 for the negative control group and n=4 for virus only and each treatment group, respectively) for virus quantitation. Animals with 25% body weight loss were euthanized and excluded from statistical analysis.

### 2.7. Virus quantitation

For the SARS-CoV-2 titration from the respiratory tract, mice were euthanized on 3 and 6 dpi, and lungs were harvested. The lung tissues were homogenized in 1 mL of DMEM media with bead-based homogenizer (Bertin Technologies). The lung supernatant samples were 10-fold serially diluted in DMEM and inoculated to Vero E6 cells in a 24-well plate. After incubation for 1 h, the inoculum was removed and covered with agar-overlay media including 0.5% (w/v) agarose (Lonza) and 2% FBS in MEM. Following incubation at 37°C for 2 days, the cells were fixed with 4% (v/v) paraformaldehyde solution, and stained with 0.2% crystal violet (Sigma). The number of plaques per well was counted and analyzed.

## 3. Results

### 3.1. *In vitro* potency of CT-P59 against Delta and its associated variants in cells

To investigate the therapeutic efficacy of CT-P59 against Delta and Epsilon variants, we first evaluated the binding affinity of CT-P59 against mutant RBDs (L452R/T478K or L452R) by using BLI (Bio-Layer interferometry). The equilibrium dissociation constant (K_D_) of CT-P59 against RBD mutant proteins of Delta or Epsilon variant was reduced by 9 ∼ 10-fold compared to that against wild-type RBD (Supplementary Table 1). Next, we carried out two types of *in vitro* assays with live or pseudotyped viruses to assess the susceptibility of Delta and its associated variants to CT-P59. CT-P59 showed reduced susceptibility against Delta, Epsilon and Kappa variants with IC_50_ of 1,237, 365.6∼499.5 and 161.5 ng/mL compared to that against wild type (6.76 ng/mL) in PRNT. Approx. 183-, 54∼74-, 24-fold reduced susceptibility against Delta, Epsilon and Kappa variants were observed compared to wild type SARS-CoV-2, respectively (Table 2 and Supplementary Fig. 1A). In addition, CT-P59 showed approx. 98-, 31-, 50-fold reduced neutralizing activity against Delta (L452R/T478K/P681R), Epsilon (S13I/W152C/L452R) and Kappa (L452R/E484Q/P681R) pseudotyped virus with an IC_50_ of 21.52 ng/mL, 6.819 ng/mL and 11.08 ng/ml compared to that of D614G pseudovirus (0.219 ng/mL), respectively (Table 2 and Supplementary Fig. 1B). Thus, we found that CT-P59 can neutralize Delta and Epsilon variants despite reduced binding affinity and antiviral ability in *in vitro* experiments. We also tested whether single mutants (L452R, T478K, and P681H) impact neutralization of CT-P59. CT-P59 was less susceptible to L452R (13.22 ng/mL), but retained its own neutralizing effect against T478K (0.213 ng/mL), and P681H (0.378 ng/mL) (Table 2 and Supplementary Fig. 1B).

**Table 1.**
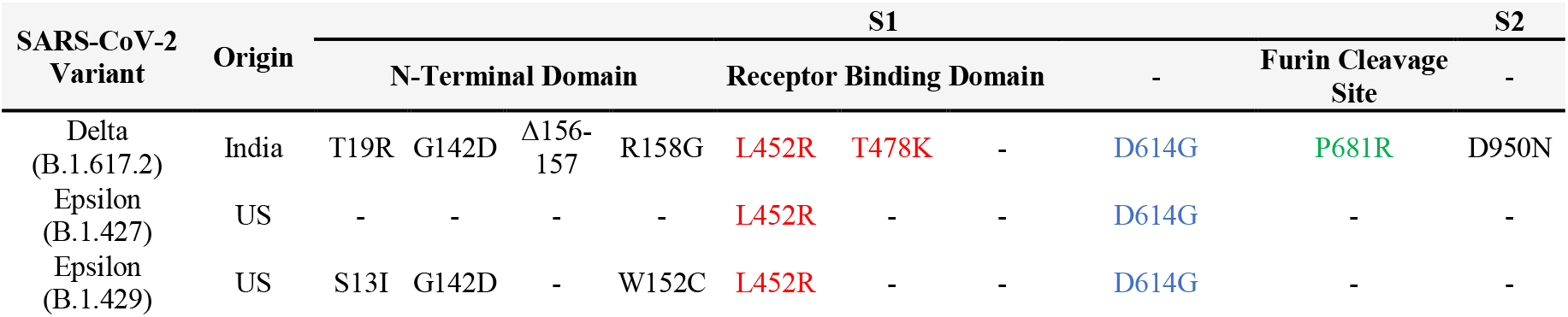
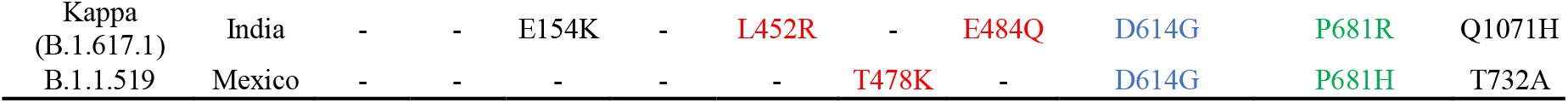
Mutations in spike protein of variants.

**Table 2.** Neutralization effect of CT-P59 against variant live viruses and pseudoviruses.

| Viruses |  | IC<sub>50</sub> (ng/mL) | Fold change (reduction) |
| --- | --- | --- | --- |
| Live virus | Delta (B.1.617.2) | 1237 (WT: 6.76) | 182.99 |
|  | Epsilon (B.1.427) | 499.5 (WT: 6.76) | 73.89 |
|  | Epsilon (B.1.429) | 365.6 (WT: 6.76) | 54.08 |
|  | Kappa (B.1.617.1) | 161.5(WT: 6.76) | 23.89 |
| Pseudovirus | Delta (L452R/T478K/P681R) | 21.52 (D614G: 0.219) | 98.26 |
|  | Epsilon (S13I/W152C/L452R) | 6.819 (D614G: 0.219) | 31.14 |
|  | Kappa (L452R/E484Q/P681R) | 11.08 (D614G: 0.219) | 50.59 |
|  | L452R | 13.22 (D614: 0.378) | 34.97 |
|  | T478K | 0.213 (D614G: 0.219) | 0.97 |
|  | P681H | 0.378 (D614: 0.378) | 1.00 |

### 3.2. *In vivo* potency of CT-P59 against Delta variant in mice

To evaluate whether CT-P59 has an ability to lower viral burden against Delta variant in *in vivo* setting, mice were treated with clinically relevant dosages in consideration of the 40 mg/kg of clinical dose of CT-P59. The dose of 80 mg/kg is considered as the equivalent exposure in clinical study (data not shown), and lower doses of 5 and 20 mg/kg were selected to investigate the lower exposure in the clinical situation.

Virus only (placebo) group were severely affected by Delta variant, showing the survival rate of 75% at 5 dpi and 0% at 6 dpi, respectively. In contrast, there was no lethality in all CT-P59 treatment groups (Fig. 1 A). CT-P59 treatment also ameliorated the weight loss of the mice infected with B.1.617.2 variants. 5 and 20 mg/kg CT-P59 treatment groups showed the delay in weight loss until 6 dpi and 40 and 80 mg/kg treatment showed statistically significant difference compared to the virus only group at 4 dpi and 5 dpi, respectively. Importantly, all CT-P59 treatment group maintained their body weight until 6 dpi, which was statistically comparable with negative control. Mean body weight loss of 5, 20, 40 and 80 mg/kg treatment groups were 14.3, 13.2, 4.6 and 5.7% at 5 dpi, respectively. In contrast, virus only group showed an average 16.3% of weight loss at 4 dpi, and eventually showed an average of 22.7% decrease for three mice while a single euthanized mouse excluded from statistical analysis at 5 dpi (Fig. 1 B).

**Figure 1.**
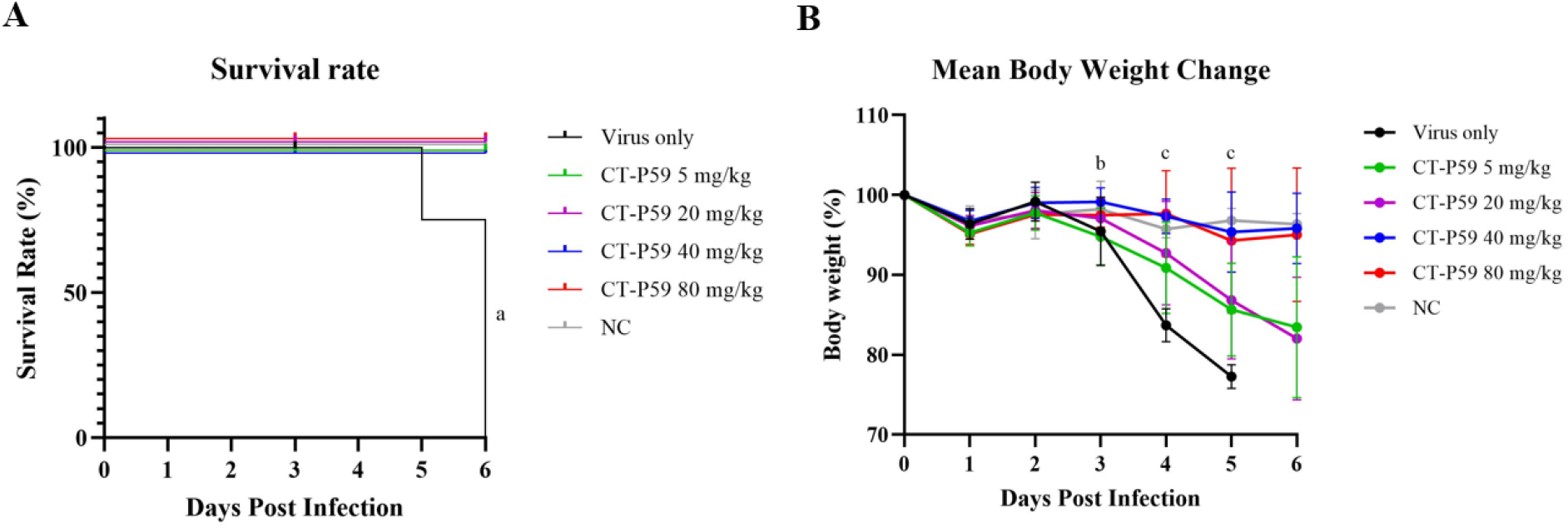
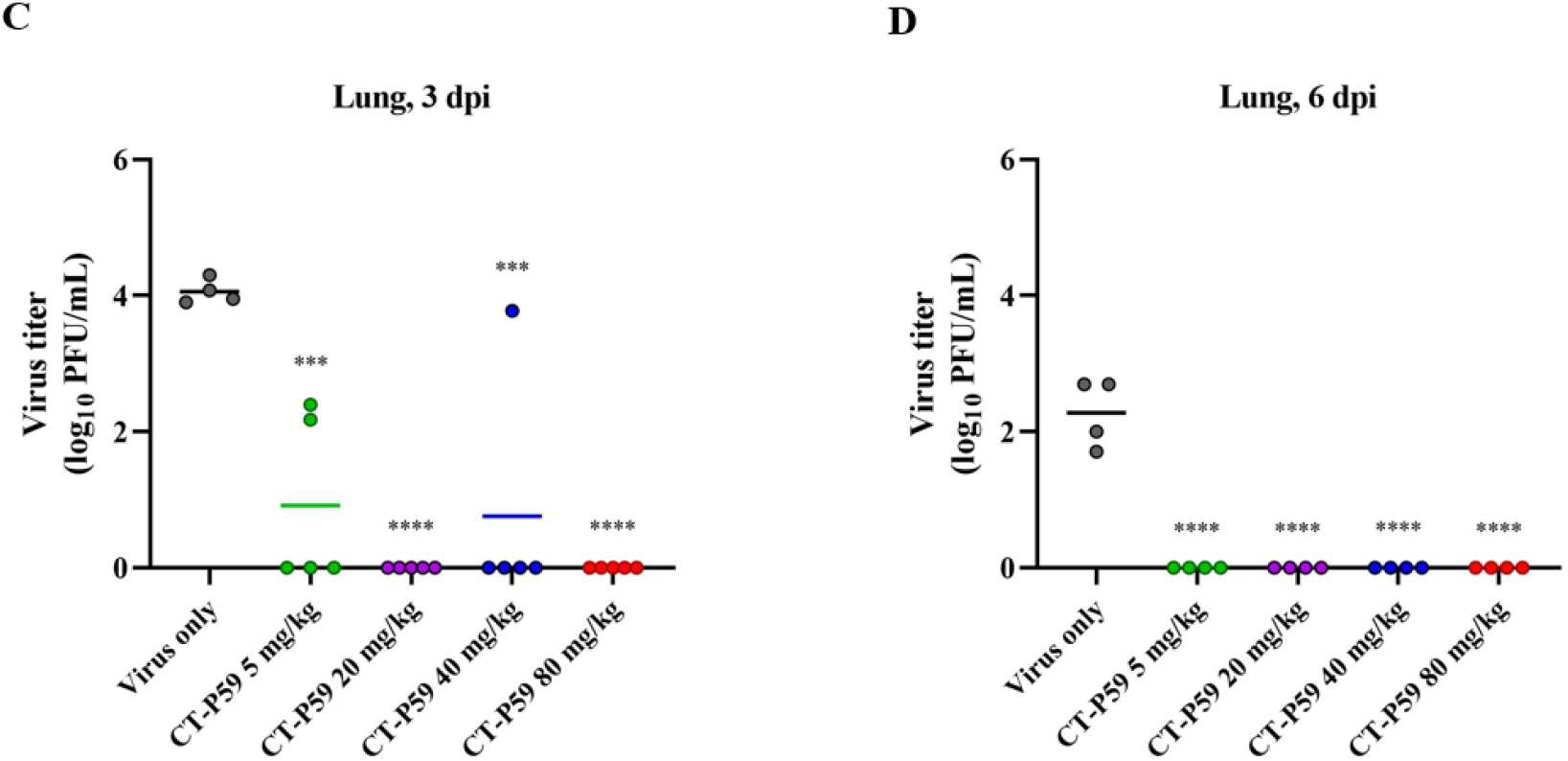
*In vivo* efficacy of clinical relevant dose of CT-P59 against Delta variant in mouse model. ACE2 mice were challenged with B.1.617.2 variants. Negative control (NC) indicates naïve mice with virus. Vehicle and 5, 20, 40 and 80 mg/kg of CT-P59 were administered intraperitoneally 8 h after virus inoculation. Body weights and survival rates were monitored daily (A, B). 25% or higher of weight loss was considered to be dead. Five and four animals each were euthanized for virus titration at 3 dpi and 6 dpi, respectively. The virus titers from lung tissue was measured (C, D) using plaque assay. Alphabets and asterisks indicates statistical significance between the virus only and each treatment group as determined by one-way ANOVA followed by a Dunnett’s post-hos test. *a* denotes statistically significant difference (p<0.05) between virus only and each treatment group (5, 20, 40 and 80 mg/kg). *b* denotes statistically significant difference (p<0.05) between virus only and 40 mg/kg. *c* denotes statistically significant difference (p<0.05) between virus only and each treatment group (40 and 80 mg/kg). *** indicates *P* < 0.001, and **** indicates *P* < 0.0001 between virus only and each treatment group.

To examine viral loads in the lower respiratory tract, we performed plaque assay with viruses from lung tissues. CT-P59 significantly abrogated the virus replication in lungs. In the virus only group, the mean virus titer in lung tissues reached 4.1 log_10_ PFU/mL at 3 dpi and declined to 2.3 log_10_ PFU/mL at 6 dpi. In contrast, all CT-P59 treatment showed substantial reduction in viral titers at 3 dpi and 6 dpi. At 3 dpi, infectious viral titers were 3.1 and 3.3 log reduced in 5 and 40 mg/kg groups, respectively, while complete reduction was observed in 20 and 80 mg/kg groups. At 6 dpi, no viral titers were detected in all CT P59 treatment (Fig 1 C and D).

## 4. Discussion

Here, we tested and confirmed *in vitro* neutralizing ability of CT-P59 against Delta, Epsilon, and Kappa variants in two different cell-based assays which are generally used to evaluate the antiviral effect of therapeutic agents. In general, it’s said that PRNT is a gold standard method to assess antivirals, since it is a cell-based assay with authentic viruses to see all the step in viral life cycle from viral entry to release, and re-infection [20]. Pseudovirus assay is broadly used since it is more convenient and known to correlate with PRNT. The advantages over PRNT are to generate mutants you want, simply detect the antiviral activities by measuring luciferase and GFP reporter, and more accessible than Bio-Safety Level 3 Laboratory. However, it’s limited to determine the *in vivo* antiviral efficacy of therapeutics since IC_50_ calculated from cell-based assay indicates its antiviral ability and it’s not reflect the effectiveness of treatment dosage and *in vivo* environments.

Like CT-P59, Sotrovimab is a monoclonal antibody which obtained the Emergency Use Authorization for the treatment of COVID-19, demonstrating the effectiveness of 500 mg dosage in clinical trial with 100 ng/mL of IC_50_ against wild type virus [21]. Also, Sotrovimab retains susceptibility against currently circulating variants such as Alpha, Beta, Gamma, Epsilon, and Kappa, in contrast to Bamlanivimab or Etesevimab [22, 23]. Regdanviamb is very potent antibody having 6 ng/mL of IC_50_ against wild type virus, which is 16.7-fold stronger than Sotrovimab [19]. Moreover, the clinical dosage of Regdanviamb come to 2,400 mg for 60 kg adult, which is 4.8-fold higher than Sotrovimab. The therapeutic ability of Regdanvimab could come to at least 80-fold (16.7 × 4.8) over that of Sotrovimab.

In accordance with *in vitro* and *in vivo* potency against Beta and Gamma variants [17, 18], CT-P59 showed reduced viral loads and clinical symptoms in Delta-infected mice. Recently, Chen *et al*., showed that a monoclonal antibody with partial loss of *in vitro* neutralizing activity confers the *in vivo* protections from variant infections in mice comparably [23]. This notion was supported by our results, suggesting that clinical dosage of CT-P59 could overcome a certain level of reduction in *in vitro* antiviral activity, as long as it is not a complete loss in neutralizing activity.

In addition, the biodistribution of monoclonal antibody to target sites *i*.*e*., respiratory tracts for COVID-19 is important to translate the *in vitro* antiviral activity to the *in vivo* efficacy in terms of pharmacokinetics of patients [24]. Given that a lung to serum concentration ratio is 15% in development of Casirivimab/Imdevimab, and Sotrovimab for the treatment of COVID-19, lung concentrations during the treatment are predicted with the *in vitro* EC_90_ to provide the rationale for clinical dosing recommendations [25-27]. CT-P59 is predicted to achieve sufficient lung epithelial lining fluid concentrations above the IC_90_ values against Delta and its associated variants during the treatment period. Hence, this biodistribution of CT-P59 supports that expected sufficient concentration in lung could compensate for reduced activity of CT-P59 against Delta and its associated variants in real world. Importantly, the finding that CT-P59 ameliorated clinical symptoms in Delta-infected mice [28] supports effectiveness of CT-P59 in patients by protection from progress into severe COVID-19. To obtain conclusive correlation between animal and human therapeutic effectiveness, it needs to be investigated in patients infected with the delta variant.

## Conclusion

In accord with previous studies with Beta and Gamma variants, this study revealed that CT-P59 retained reduced activity to Delta, Epsilon, and Kappa variants in cells, and showed *in vivo* substantial protection in Delta variant-infected mice, suggesting that the use of CT-P59 could be a curable option to COVID-19 patients infected with these variants and potential variants including identical mutation sites.

## Acknowledgements

This study was supported by a Korea National Institute of Health fund (2021-NI-007-00, 2020-ER5311-00).

## Supplementary Material

**Supplementary Table 1.** Binding affinity between CT-P59 and variant RBD proteins.

| Ligand<sup>1</sup> | Analyte | K<sub>D</sub> (M) | Fold change (reduction) |
| --- | --- | --- | --- |
| L452R/T478K | CT-P59 | 5.41E-10 (WT: 5.56E-11) | 10 |
| L452R | CT-P59 | 4.82E-10 (WT: 5.10E-11) | 9 |

<sup>1</sup>RBD (wild-type) was prepared in-house and mutant RBDs were purchased from Sino Biological.

**Supplementary figure 1.**
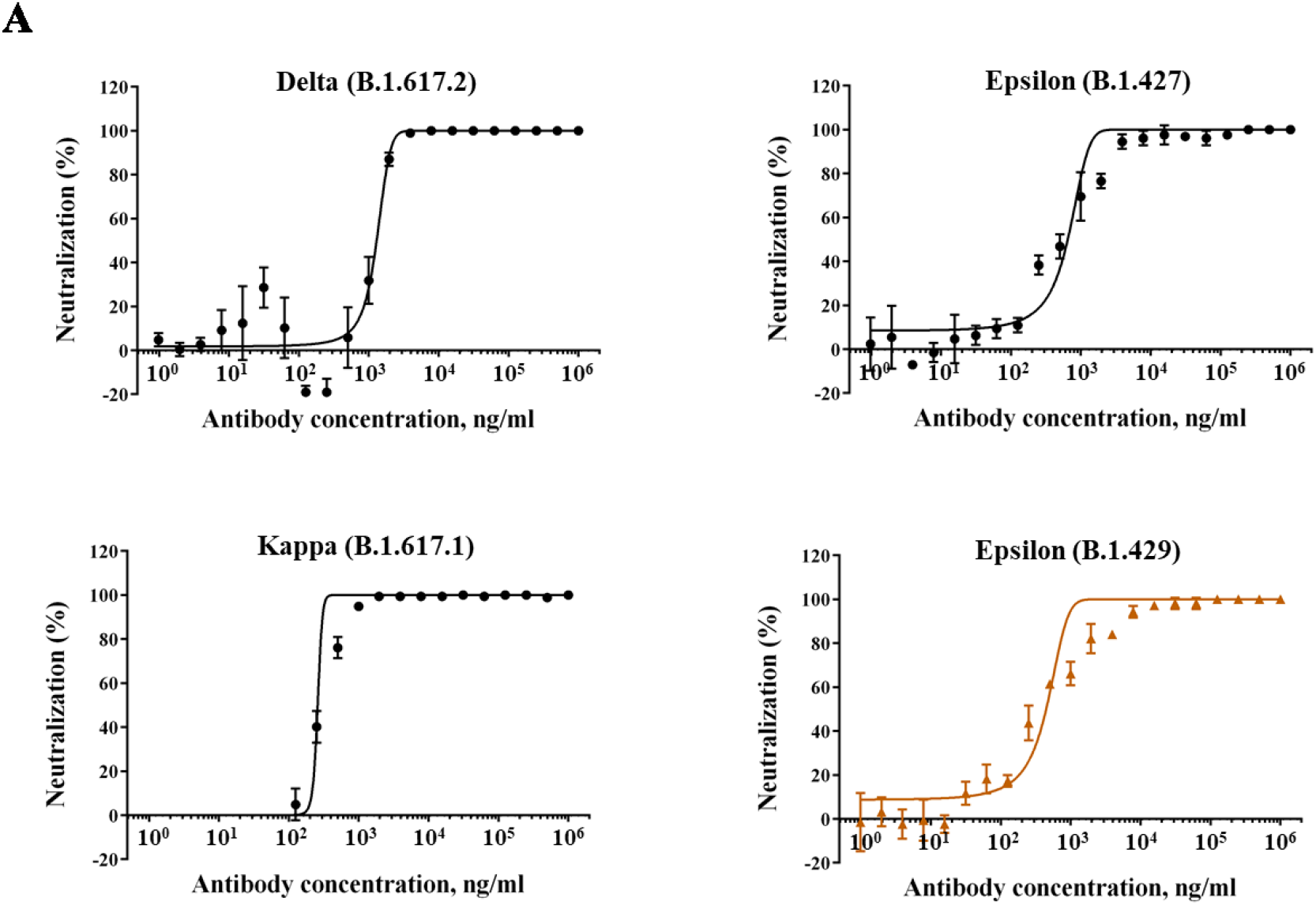
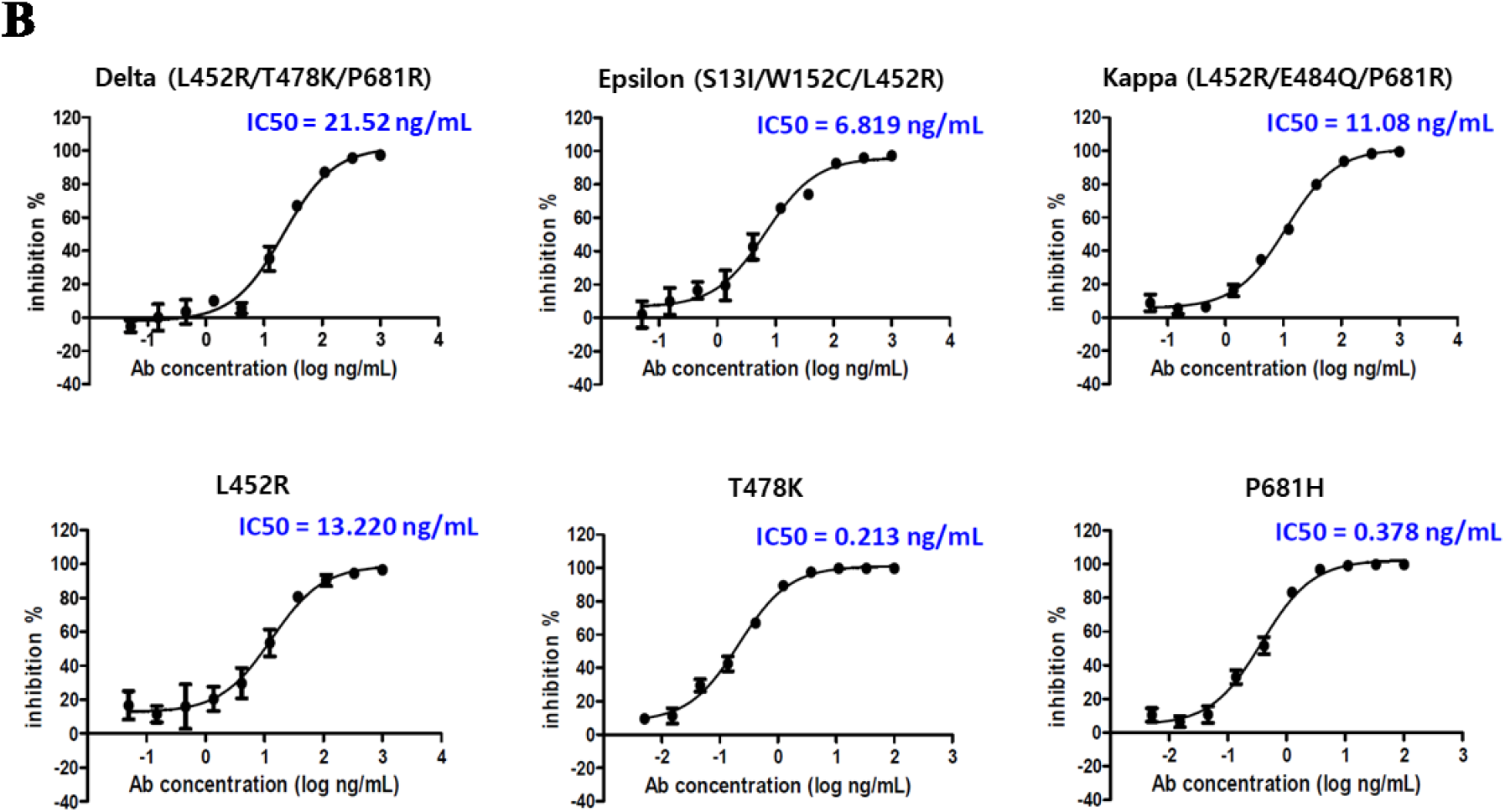
*In vitro* susceptibility test of CT-P59 against Delta, Epsilon, and Kappa variants. Two fold-serially diluted CT-P59 were incubated with various SARS-CoV-2 variants such as Delta (B.1.617.2), Epsilon (B.1.427/429), Kappa (B.1.617.1). Each antibody-virus mixture was added to VeroE6 cells. After 2–3 days of incubation, the neutralization activity was evaluated by counting plaques (A). Serially diluted CT-P59 was premixed with indicated variant pseudoviruses followed by inoculation to ACE2-expressing HEK293T cells. Luciferase activity was measured to calculate %Neutralization of CT-P59.

## References

[1] F. Campbell, B. Archer, H. Laurenson-Schafer, Y. Jinnai, F. Konings, N. Batra, B. Pavlin, K. Vandemaele, M.D. Van Kerkhove, T. Jombart, O. Morgan, O. le Polain de Waroux, Increased transmissibility and global spread of SARS-CoV-2 variants of concern as at June 2021, Euro Surveill, 26 2021.

[2] V.P. Sarah Cherian, Santosh Jadhav, Pragya Yadav, Nivedita Gupta, Mousmi Das, Partha Rakshit, Sujeet Singh, Priya Abraham, Samiran Panda, NIC team, Convergent evolution of SARS-CoV-2 spike mutations, L452R, E484Q and P681R, in the second wave of COVID-19 in Maharashtra, India, BioRxiv, (2021).

[3] P.H. England, Variants: distribution of case data, (2021).

[4] C.f.D.C.a. Prevention, Variant Proportions, (2021).

[5] D. Planas, D. Veyer, A. Baidaliuk, I. Staropoli, F. Guivel-Benhassine, M.M. Rajah, C. Planchais, F. Porrot, N. Robillard, J. Puech, M. Prot, F. Gallais, P. Gantner, A. Velay, J. Le Guen, N. Kassis-Chikhani, D. Edriss, L. Belec, A. Seve, L. Courtellemont, H. Pere, L. Hocqueloux, S. Fafi-Kremer, T. Prazuck, H. Mouquet, T. Bruel, E. Simon-Loriere, F.A. Rey, O. Schwartz, Reduced sensitivity of SARS-CoV-2 variant Delta to antibody neutralization, Nature, (2021).

[6] D.V. Delphine Planas, Artem Baidaliuk, Isabelle Staropoli, Florence Guivel-Benhassine, Maaran Michael Rajah, Cyril Planchais, Françoise Porrot, Nicolas Robillard, Julien Puech, Matthieu Prot, Floriane Gallais, Pierre Gantner, Aurélie Velay, Julien Le Guen, Najibi Kassis-Chikhani, Dhiaeddine Edriss, Laurent Belec, Aymeric Seve, Hélène Péré, Laura Courtellemont, Laurent Hocqueloux, Samira Fafi-Kremer, Thierry Prazuck, Hugo Mouquet, Timothée Bruel, Etienne Simon-Lorière, Felix Rey, Olivier Schwartz, Reduced sensitivity of infectious SARS-CoV-2 variant B.1.617.2 to monoclonal antibodies and sera from convalescent and vaccinated individuals, bioRxiv, (2021).

[7] T.N. Starr, A.J. Greaney, A. Addetia, W.W. Hannon, M.C. Choudhary, A.S. Dingens, J.Z. Li, J.D. Bloom, Prospective mapping of viral mutations that escape antibodies used to treat COVID-19, Science, 371 (2021) 850–854.

[8] D. Planas, T. Bruel, L. Grzelak, F. Guivel-Benhassine, I. Staropoli, F. Porrot, C. Planchais, J. Buchrieser, M.M. Rajah, E. Bishop, M. Albert, F. Donati, M. Prot, S. Behillil, V. Enouf, M. Maquart, M. Smati-Lafarge, E. Varon, F. Schortgen, L. Yahyaoui, M. Gonzalez, J. De Seze, H. Pere, D. Veyer Seve, E. Simon-Loriere, S. Fafi-Kremer, K. Stefic, H. Mouquet, L. Hocqueloux, S. van der Werf, T. Prazuck, O. Schwartz, Sensitivity of infectious SARS-CoV-2 B.1.1.7 and B.1.351 variants to neutralizing antibodies, Nat Med, 27 (2021) 917–924.

[9] S.M. Pragya D. Yadav, Anita M Shete, Dimpal A Nyayanit, Nivedita Gupta, Deepak Y. Patil, Gajanan N. Sapkal, Varsha Potdar, Manoj Kadam, Abhimanyu Kumar, Sanjay Kumar, Deepak Suryavanshi, Chandrashekhar S. Mote, Priya Abraham, Samiran Panda, Balram Bhargava, SARS CoV-2 variant B.1.617.1 is highly pathogenic in hamsters than B.1 variant, bioRxiv, (2021).

[10] M. McCallum, J. Bassi, A. De Marco, A. Chen, A.C. Walls, J. Di Iulio, M.A. Tortorici, M.J. Navarro, C. Silacci-Fregni, C. Saliba, K.R. Sprouse, M. Agostini, D. Pinto, K. Culap, S. Bianchi, S. Jaconi, E. Cameroni, J.E. Bowen, S.W. Tilles, M.S. Pizzuto, S.B. Guastalla, G. Bona, A.F. Pellanda Garzoni, W.C. Van Voorhis, L.E. Rosen, G. Snell, A. Telenti, H.W. Virgin, L. Piccoli, D. Corti, D. Veesler, SARS-CoV-2 immune evasion by the B.1.427/B.1.429 variant of concern, Science, (2021).

[11] X. Deng, M.A. Garcia-Knight, M.M. Khalid, V. Servellita, C. Wang, M.K. Morris, A. Sotomayor-Gonzalez, D.R. Glasner, K.R. Reyes, A.S. Gliwa, N.P. Reddy, C. Sanchez San Martin, S. Federman, J. Cheng, J. Balcerek, J. Taylor, J.A. Streithorst, S. Miller, B. Sreekumar, P.Y. Chen, U. Schulze-Gahmen, T.Y. Taha, J.M. Hayashi, C.R. Simoneau, G.R. Kumar, S. McMahon, P.V. Lidsky, Y. Xiao, P. Hemarajata, N.M. Green, A. Espinosa, C. Kath, M. Haw, J. Bell, J.K. Hacker, C. Hanson, D.A. Wadford, C. Anaya, D. Ferguson, P.A. Frankino, H. Shivram, L.F. Lareau, S.K. Wyman, M. Ott, R. Andino, C.Y. Chiu, Transmission, infectivity, and neutralization of a spike L452R SARS-CoV-2 variant, Cell, 184 (2021) 3426–3437 e3428.

[12] M. McCallum, J. Bassi, A. Marco, A. Chen, A.C. Walls, J.D. Iulio, M.A. Tortorici, M.J. Navarro Silacci-Fregni, C. Saliba, M. Agostini, D. Pinto, K. Culap, S. Bianchi, S. Jaconi, E. Cameroni, J.E. Bowen, S.W. Tilles, M.S. Pizzuto, S.B. Guastalla, G. Bona, A.F. Pellanda, C. Garzoni, W.C. Van Voorhis, L.E. Rosen, G. Snell, A. Telenti, H.W. Virgin, L. Piccoli, D. Corti, D. Veesler, SARS-CoV-2 immune evasion by variant B.1.427/B.1.429, bioRxiv, (2021).

[13] H.N. Akatsuki Saito, Keiya Uriu, Yusuke Kosugi, Takashi Irie, Kotaro Shirakawa, Kenji Sadamasu, Izumi Kimura, Jumpei Ito, Jiaqi Wu, Seiya Ozono, Kenzo Tokunaga, Erika P Butlertanaka, Yuri L Tanaka, Ryo Shimizu, Kenta Shimizu, Takasuke Fukuhara, Ryoko Kawabata, Takemasa Sakaguchi, Isao Yoshida, Hiroyuki Asakura, Mami Nagashima, Kazuhisa Yoshimura, Yasuhiro Kazuma, Ryosuke Nomura, Yoshihito Horisawa, Akifumi Takaori-Kondo, The Genotype to Phenotype Japan (G2P-Japan) Consortium, So Nakagawa, Terumasa Ikeda, Kei Sato, SARS-CoV-2 spike P681R mutation enhances and accelerates viral fusion, bioRxiv, (2021).

[14] C.M.S. Thomas P. Peacock, Jonathan C. Brown, Niluka Goonawardane, Jie Zhou, Max Whiteley, PHE Virology Consortium, Thushan I. de Silva, Wendy S. Barclay, The SARS-CoV-2 variants associated with infections in India, B.1.617, show enhanced spike cleavage by furin, bioRxiv, (2021).

[15] B. Lubinski, T. Tang, S. Daniel, J.A. Jaimes, G.R. Whittaker, Functional evaluation of proteolytic activation for the SARS-CoV-2 variant B.1.1.7: role of the P681H mutation, bioRxiv, (2021).

[16] D.P. Maison, L.L. Ching, C.M. Shikuma, V.R. Nerurkar, Genetic Characteristics and Phylogeny of 969-bp S Gene Sequence of SARS-CoV-2 from Hawaii Reveals the Worldwide Emerging P681H Mutation, bioRxiv, (2021).

[17] D.K. Ryu, R. Song, M. Kim, Y.I. Kim, C. Kim, J.I. Kim, K.S. Kwon, A.S. Tijsma, P.M. Nuijten, C.A. van Baalen, T. Hermanus, P. Kgagudi, T. Moyo-Gwete, P.L. Moore, Y.K. Choi, S.Y. Lee, Therapeutic effect of CT-P59 against SARS-CoV-2 South African variant, Biochem Biophys Res Commun, 566 (2021) 135–140.

[18] B.K. Dong-Kyun Ryu, Sun-Je Woo, Min-Ho Lee, Aloys SL Tijsma, Hanmi Noh, Jong-In Kim, Ji-Min Seo, Cheolmin Kim, Minsoo Kim, Eunji Yang, Gippeum Lim, Seong-Gyu Kim, Su-Kyeong Eo, Jung-ah Choi, Sang-Seok Oh, Patricia M Nuijten, Manki Song, Hyo-Young Chung, Carel A van Baalen, Ki-Sung Kwon, Soo-Young Lee, Therapeutic efficacy of CT-P59 against P.1 variant of SARS-CoV-2, BioRxiv, (2021).

[19] C. Kim, D.K. Ryu, J. Lee, Y.I. Kim, J.M. Seo, Y.G. Kim, J.H. Jeong, M. Kim, J.I. Kim, P. Kim, J.S. Bae, E.Y. Shim, M.S. Lee, M.S. Kim, H. Noh, G.S. Park, J.S. Park, D. Son, Y. An, J.N. Lee, K.S. Kwon, J.Y. Lee, H. Lee, J.S. Yang, K.C. Kim, S.S. Kim, H.M. Woo, J.W. Kim, M.S. Park, K.M. Yu, S.M. Kim, E.H. Kim, S.J. Park, S.T. Jeong, C.H. Yu, Y. Song, S.H. Gu, H. Oh, B.S. Koo, J.J. Hong, C.M. Ryu, W.B. Park, M.D. Oh, Y.K. Choi, S.Y. Lee, A therapeutic neutralizing antibody targeting receptor binding domain of SARS-CoV-2 spike protein, Nat Commun, 12 (2021) 288.

[20] K.R. Bewley, N.S. Coombes, L. Gagnon, L. McInroy, N. Baker, I. Shaik, J.R. St-Jean, N. St-Amant, K.R. Buttigieg, H.E. Humphries, K.J. Godwin, E. Brunt, L. Allen, S. Leung, P.J. Brown, E.J. Penn, K. Thomas, G. Kulnis, B. Hallis, M. Carroll, S. Funnell, S. Charlton, Quantification of SARS-CoV-2 neutralizing antibody by wild-type plaque reduction neutralization, microneutralization and pseudotyped virus neutralization assays, Nat Protoc, 16 (2021) 3114–3140.

[21] C.H.-D. Andrea L. Cathcart, Florian A. Lempp, Daphne Ma, Michael A. Schmid, Maria L. Agostini, Barbara Guarino, Julia Di iulio, Laura E. Rosen, Heather Tucker, Joshua Dillen, Sambhavi Subramanian, Barbara Sloan, Siro Bianchi, Dora Pinto, Christian Saliba, Jason A Wojcechowskyj, Julia Noack, Jiayi Zhou, Hannah Kaiser, Arthur Chase, Martin Montiel-Ruiz, Exequiel Dellota Jr., Arnold Park, Roberto Spreafico, Anna Sahakyan, Elvin J. Lauron, Nadine Czudnochowski, Elisabetta Cameroni, Sarah Ledoux, Adam Werts, Christophe Colas, Leah Soriaga, Amalio Telenti, Lisa A. Purcell, Seungmin Hwang, Gyorgy Snell, Herbert W. Virgin, Davide Corti, Christy M. Hebner, The dual function monoclonal antibodies VIR-7831 and VIR-7832 demonstrate potent in vitro and in vivo activity against SARS-CoV-2, (2021).

[22] T.N. Starr, A.J. Greaney, A.S. Dingens, J.D. Bloom, Complete map of SARS-CoV-2 RBD mutations that escape the monoclonal antibody LY-CoV555 and its cocktail with LY-CoV016, Cell Rep Med, 2 (2021) 100255.

[23] R.E. Chen, E.S. Winkler, J.B. Case, I.D. Aziati, T.L. Bricker, A. Joshi, T.L. Darling, B. Ying, J.M. Errico, S. Shrihari, L.A. VanBlargan, X. Xie, P. Gilchuk, S.J. Zost, L. Droit, Z. Liu, S. Stumpf, Wang, S.A. Handley, W.B. Stine, Jr., P.Y. Shi, M.E. Davis-Gardner, M.S. Suthar, M.G. Knight, R. Andino, C.Y. Chiu, A.H. Ellebedy, D.H. Fremont, S.P.J. Whelan, J.E. Crowe, Jr., L. Purcell, D. Corti, A.C.M. Boon, M.S. Diamond, In vivo monoclonal antibody efficacy against SARS-CoV-2 variant strains, Nature, (2021).

[24] R. Jia, H. Wang, How Could In Vitro Antiviral Activity Be Applied to Optimize the Dosing Regimens of Candidates for the Treatment of Severe Acute Respiratory Syndrome Coronavirus 2 (SARS-CoV-2)?, Clin Infect Dis, 73 (2021) 352–353.

[25] DA, Fact sheet for health care providers ememgency use authorization (EUA) of casirivimab and imdevimab, (2020).

[26] FDA, Fact sheet for healthcare providers emegency use authorization (EUA) of sotrovimab, (2021).

[27] D.K. Shah, A.M. Betts, Antibody biodistribution coefficients: inferring tissue concentrations of monoclonal antibodies based on the plasma concentrations in several preclinical species and human, MAbs, 5 (2013) 297–305.

[28] M.D. Johansen, A. Irving, X. Montagutelli, M.D. Tate, I. Rudloff, M.F. Nold, N.G. Hansbro, R.Y. Kim, C. Donovan, G. Liu, A. Faiz, K.R. Short, J.G. Lyons, G.W. McCaughan, M.D. Gorrell, A. Cole, C. Moreno, D. Couteur, D. Hesselson, J. Triccas, G.G. Neely, J.R. Gamble, S.J. Simpson, B.M. Saunders, B.G. Oliver, W.J. Britton, P.A. Wark, C.A. Nold-Petry, P.M. Hansbro, Animal and translational models of SARS-CoV-2 infection and COVID-19, Mucosal Immunol, 13 (2020) 877–891.

